# Impact of antimicrobial treatment for acute otitis media on carriage dynamics of penicillin-susceptible and penicillin–non-susceptible *Streptococcus pneumoniae*: secondary analysis of a randomized, double-blind, placebo-controlled trial

**DOI:** 10.1101/236752

**Authors:** Joseph A. Lewnard, Paula A. Tähtinen, Miia K. Laine, Laura Lindholm, Jari Jalava, Pentti Huovinen, Marc Lipsitch, Aino Ruohola

**Affiliations:** Center for Communicable Disease Dynamics, Department of Epidemiology, Harvard TH Chan School of Public Health, Boston, MA 02115; Department of Paediatrics and Adolescent Medicine, Turku University Hospital, Turku, Finland; Department of Paediatrics and Adolescent Medicine, University of Turku, Turku, Finland; Department of Clinical Microbiology, Turku University Hospital, Turku, Finland; Expert Microbiology Unit, National Institute for Health and Welfare, Helsinki, Finland; Infectious Disease Control and Vaccinations Unit, National Institute for Health and Welfare, Helsinki, Finland; Department of Medical Microbiology and Immunology, University of Turku, Turku, Finland

## Abstract

**Background:** Concerns that antimicrobial treatment may foster selection and transmission of resistant bacterial lineages have led to conflicting guidelines for clinical management of common non-severe infections. However, the impact of antimicrobial treatment on colonization dynamics is poorly understood. We used data from a previously-conducted trial of amoxicillin-clavulanate therapy for acute otitis media (AOM) to understand how antimicrobial treatment impacts the acquisition and clearance of *Streptococcus pneumoniae* lineages with varying susceptibility to penicillin.

**Methods and findings:** We measured impacts of antimicrobial treatment on nasopharyngeal carriage of penicillin-susceptible *S. pneumoniae* (PSSP) and penicillin–non-susceptible *S. pneumoniae* (PNSP) lineages at end-of-treatment and 15d, 30d, and 60d after treatment in a previously-conducted randomized, double-blind, placebo-controlled trial. Analyses were not specified in the original protocol. Among children 6-35 months of age with stringently-defined AOM, 162 were assigned amoxicillin-clavulanate, and 160 were assigned placebo. Children who did not show clinical improvement received open-label antimicrobial rescue treatment with amoxicillin-clavulanate irrespective of the randomized treatment assignment, to which both patients and physicians were blinded. The intention-to-treat populations of the intervention and placebo arms thus received care resembling immediate antimicrobial therapy and watchful waiting, respectively. Immediate amoxicillin-clavulanate reduced PSSP carriage prevalence by 88% (95%CI: 76-96%) at end-of-treatment and by 27% (–3-49%) after 60d, but did not measurably alter PNSP carriage prevalence throughout follow-up. By end-of-treatment, 7% of children who carried PSSP at enrollment remained colonized in the amoxicillin-clavulanate arm, compared to 61% of PSSP carriers who received placebo; differences in carriage prevalence persisted at least 60d after treatment among children who carried PSSP at enrollment. Among children not carrying pneumococci at enrollment, amoxicillin-clavulanate reduced PSSP acquisition by >80% over 15d. Among children who carried PNSP at enrollment, no differences in carriage prevalence of *S. pneumoniae*, PSSP, or PNSP were detected at follow-up visits.

**Conclusions:** In a setting with low PNSP prevalence, antimicrobial therapy for AOM conferred a selective impact on colonizing *S. pneumoniae* by accelerating clearance, and delaying acquisition, of penicillin-susceptible lineages. Absolute risk of carrying PNSP was unaffected by treatment (ClinicalTrials.gov: NCT00299455; Funding: NIH/NIGMS).

## INTRODUCTION

Acute otitis media (AOM) is the leading cause of healthcare visits and antimicrobial prescribing among children in high-income countries [1]. Although many infections are self-limiting, antimicrobial therapy accelerates the resolution of otoscopic signs and symptoms [2,3]. Because AOM is prevalent among children, routinely treating infections with antimicrobial therapy may exert considerable selective pressure in favor of drug resistance. This concern has led to conflicting views and guidelines about the optimal clinical management of AOM [4], with certain countries recommending “watchful waiting” as a strategy to limit unnecessary antimicrobial prescribing for infections that may resolve spontaneously [5,6].

Limited evidence of how treating AOM with antimicrobial drugs impacts selection of resistant bacteria has contributed to uncertainty about the extent to which resistance risks may offset clinical benefits of treatment [4]. Associations between previous antimicrobial treatment and carriage of resistant bacteria have been reported in observational studies [7,8]. However, endpoints of drug-susceptible and drug-resistant bacterial carriage have been measured in few trials, and incompatible reporting of such endpoints across individual trials has made outcomes difficult to interpret [3,9–11]. Effects of antimicrobial treatment on the clearance, replacement, or acquisition of bacterial strains, and the persistence of these effects over time, have not been well-quantified. Because such data are integral to the parameterization of mathematical models of selection and transmission dynamics [12], this gap has curtailed our ability to assess the impacts of alternative treatment protocols on the spread of drug-resistant bacteria [5,6], and to evaluate the population-level benefits of interventions aiming to combat antimicrobial resistance [13,14].

A previously-conducted clinical trial of immediate amoxicillin clavulanate for AOM provided an opportunity to assess how antimicrobial treatment impacts nasopharyngeal carriage of *Streptococcus pneumoniae* lineages with varying susceptibility to penicillin [2]. To improve the evidence base for treatment guidelines and quantitative modeling of transmission, we revisited data from this study, measuring treatment effects on carriage up to two months after treatment.

## METHODS

### Study design and patients

Nasopharyngeal carriage of *S. pneumoniae* was assessed as a secondary outcome in a randomized, double-blind, placebo-controlled trial of amoxicillin-clavulanate for the treatment of stringently-defined AOM at Turku University Hospital in Turku, Finland. Trial design, primary clinical endpoints, adverse events, and the protocol of the original have been reported previously [2]. Briefly, children 6 to 35 months old seeking care were screened for AOM diagnosis, defined as fulfilling the following three criteria: presence of middle ear fluid, acute inflammatory signs in the tympanic membrane, and acute symptoms such as fever, ear pain, or respiratory symptoms. Enrollment ran from March 2006 to December 2008. In this manuscript, we present analyses of secondary endpoints, were not planned in the protocol for the original study.

Upon enrollment, children were randomized to immediate antimicrobial therapy or placebo, and attended scheduled follow-up visits with study physicians. Rescue therapy with open-label antimicrobial drugs was initiated if children’s condition did not improve or worsened during treatment, based on parental report and clinical assessment of symptoms and otoscopic signs. In this sense, the intention-to-treat populations of the treatment and placebo arms were blinded to prescribing strategies that resembled immediate antimicrobial therapy and watchful waiting, respectively. Children receiving rescue treatment were thus retained in the placebo study arm for analyses to determine the effect of immediate antimicrobial prescribing.

Nasopharyngeal swabbing occurred in-clinic at enrollment, at scheduled visits, and when children sought care for new episodes. We analyzed carriage outcomes for scheduled visits occurring at end-of-treatment (day 8) and 15, 30, and 60 days after treatment. So that follow-up visits occurred with equal time after end-of-treatment for all children, study days were counted from the enrollment visit for children who did not require rescue treatment, and from the day of initiating rescue treatment for children who required such therapy. Susceptibility of isolates to penicillin was determined using standard breakpoints, as detailed below [15].

### Ethics

Parents provided written informed consent on behalf of children for all study procedures. The ethics committee of the Hospital District of Southwest Finland Ethical approved the original study. Secondary analyses were exempted by the institutional review board at Harvard TH Chan School of Public Health.

### Randomization and masking

Patients were enrolled by study physicians and randomly assigned (1:1) amoxicillin-clavulanate or placebo via computer-generated allocation numbers, which were provided over the telephone by nurses in the Pediatric Infectious Disease Ward of Turku University Hospital. Patients, study physicians, and laboratory personnel were blinded to the randomization codes for the duration of the trial. The allocation list was generated as a block of ten and kept by the hospital pharmacy. Pharmacists concealed assignments by labeling identical study drug containers with allocation numbers. Placebo treatments (capsules containing lactose monohydrate, 640mg) were identical to the active treatment in appearance, taste, and smell. The biostatistician performing analyses presented in this manuscript was not blinded to treatment assignments.

### Procedures

Eligible patients were randomized to amoxicillin-clavulanate (40mg of amoxicillin plus 5.7mg of clavulanate per kg of body weight daily, divided into two doses) or placebo for 7 days. Study physicians encouraged the use of analgesic and antipyretic agents, and allowed the use of decongestant nose drops and sprays. To enable comparisons of immediate versus delayed antimicrobial treatment, the main rescue treatment was open-label amoxicillin-clavulanate 40/5.7mg per kg/day, divided into two doses, for 7 days. However, study physicians could also prescribe high-dose amoxicillin-clavulanate or intramuscular ceftriaxone as rescue treatment based on clinical assessment, consistent with Finnish treatment guidelines.

Nasopharyngeal specimens were collected with dacron swabs (Copan diagnostics, Corona, CA USA), and plated on selective agar immediately after the study visit to determine pneumococcal carriage. Plates were incubated in 5% CO_2_ at ≥35°C and examined at 18-24h and 36-48h. Penicillin susceptibility was determined via a two-step approach. Samples were first screened for reduced susceptibility using oxacillin disks. For samples with a decreased (<20mm) zone of inhibition, minimal inhibitory concentrations (MICs) were determined using penicillin G E-tests (bioMérieux, Cambridge, MA, USA). All samples with a zone of exclusion ≥20mm, or a zone of exclusion <20mm and MIC ≤0.064µg/mL, were classed as penicillin-susceptible. Intermediate resistance was defined as a zone of exclusion <20mm and MIC >0.064µg/mL or ≤2µg/mL, and resistance was defined as a zone of exclusion <20mm and MIC >2µg/mL. A susceptible control strain (ATCC 49619) was tested together with clinical isolates for validation.

### Outcomes

The endpoints of interest were carriage of any pneumococci, carriage of penicillin-susceptible *S. pneumoniae* (PSSP), and carriage of penicillin–non-susceptible *S. pneumoniae* (PNSP), defined as an intermediately-resistant or resistant phenotype, at 8, 15, 30, and 60 days after the start of treatment with the study drug or rescue therapy.

### Statistical analysis

#### Definition of treatment effects

Children were analyzed according to the study arms to which they were randomized in an intention-to-treat framework. We calculated the risk ratio (RR) of carriage endpoints in the two arms via the prevalence ratio of any pneumococcal carriage, of PSSP carriage, and of PNSP carriage. We measured the impact of immediate antimicrobial prescribing against each colonization endpoint at follow-up visits as (1–*RR*)×100%. To understand effects of treatment on acquisition and clearance of *S. pneumoniae*, we also performed subgroup analyses comparing prevalence of any pneumococcal carriage, of PSSP carriage, and of PNSP carriage among children who, at enrollment, were not colonized, or carried PSSP or PNSP.

To determine the selective impact of antimicrobial treatment on pneumococci carried by children treated for AOM, we next assessed the susceptibility of strains carried by children who had been assigned to immediate amoxicillin-clavulanate or placebo [11]. We measured the conditional risk ratio (cRR) for strains carried at follow-up visits to show reduced susceptibility as the ratio, between the study arms, of the proportions of pneumococcal isolates classed as PNSP, out of all pneumococcal isolates tested.

Last, we calculated risk differences to measure the prevalence of carriage that could be attributed to, or averted by, immediate antimicrobial prescribing. For each scheduled visit, we calculated differences in pneumococcal carriage prevalence—including all pneumococci, PSSP, and PNSP—between the two study arms. We again performed subgroup analyses according to carriage status at enrollment.

#### Measurement of prevalence and statistical testing

We conducted statistical inference over 100,000 bootstrap-resampled datasets to measure medians and 95% confidence intervals (CIs) around prevalence, risk ratio, and risk difference estimates. As detailed in the supplemental text (**Text S1**) the bootstrap approach allowed us to generate unbiased estimates of PSSP and PNSP carriage prevalence in the presence of missing observations that would otherwise bias estimates.

## RESULTS

### Enrollment and sample processing

Of 162 patients initially randomized to amoxicillin-clavulanate and 160 initially randomized to placebo, two children in the treatment arm and three children in the placebo arm left the study prior to the end-of-treatment visit (**Figure 1**). Rescue therapy was initiated in 11 (7%) children randomized to amoxicillin-clavulanate, compared to 53 (33%) children randomized to placebo.

**Figure 1:**
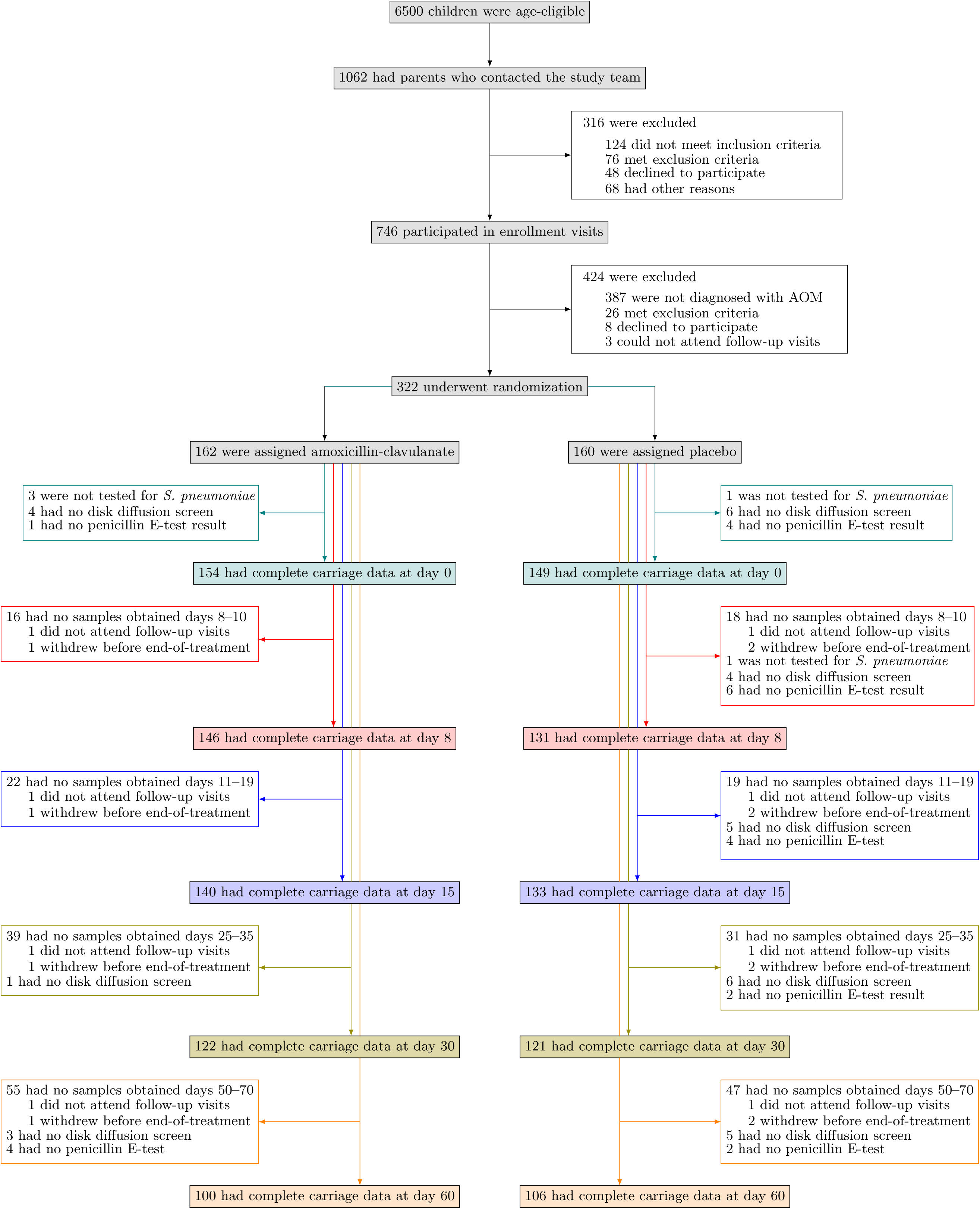
Trial outline. We illustrate the recruitment, enrollment, and randomization of patients together with the completion of follow-up visits as scheduled and processing of nasopharyngeal specimens. Samples were not obtained if children did not attend a visit within the specified time windows of scheduled visits at days 8, 15, 30, and 60, or if children or parents refused the nasopharyngeal swab.

At enrollment, 193 of 318 (61%) children carried *S. pneumoniae* (**Table 1**); the most frequently-carried serotypes were 6B (*n=*60), 19F (*n=*56), 14 (*n=*35), 23F (*n=*27), 6A (*n=*15), 6C (*n=*11), and 35B (*n=*10). In total, pneumococcal carriage was determined at 675 of 810 (83%) scheduled visits among children assigned amoxicillin-clavulanate, and at 683 of 800 (85%) scheduled visits among children assigned placebo (**Figure 1**). Resistance determinations were available for 662 (98%) and 640 (94%) pneumococcal isolates in these arms, respectively. Baseline characteristics of children with complete follow-up data to day 60 did not differ from baseline characteristics of children missing data from any scheduled visit (**Table S1**).

**Table 1:** Baseline characteristics of patients.

| Characteristic | All children |  | Children carrying *S. pneumoniae* at enrollment |  | Children not colonized by *S. pneumoniae* at enrollment |  |
| --- | --- | --- | --- | --- | --- | --- |
|  | Amox.-Clav.<br>(N=162) | Placebo<br>(N=160) | Amox.-Clav.<br>(N=102) | Placebo<br>(N=91) | Amox.-Clav.<br>(N=57) | Placebo<br>(N=68) |
| Age (mo.) |  |  |  |  |  |  |
| Mean (±SD) | 15.6 (±7.3) | 16.0 (±7.9) | 15.7 (±7.3) | 16.4 (±8.2) | 15.5 (±7.4) | 15.5 (±7.5) |
| Male child |  |  |  |  |  |  |
| N (%) | 92 (57) | 91 (57) | 59 (58) | 46 (51) | 31 (54) | 45 (66) |
| Received antimicrobial drugs within preceding month |  |  |  |  |  |  |
| N (%) | 37 (23) | 34 (21) | 21 (21) | 14 (15) | 15 (26) | 20 (29) |
| Number of siblings |  |  |  |  |  |  |
| 0, N (%) | 73 (45) | 66 (41) | 46 (45) | 34 (37) | 24 (42) | 31 (46) |
| 1–2, N (%) | 79 (49) | 89 (56) | 51 (50) | 53 (58) | 28 (49) | 36 (53) |
| 3 or more, N (%) | 10 (6) | 5 (3) | 5 (5) | 4 (4) | 5 (9) | 1 (1) |
| Daycare attendance |  |  |  |  |  |  |
| N (%) | 87 (54) | 87 (54) | 63 (47) | 49 (34) | 23 (30) | 37 (43) |
| Parental smoking<sup>1</sup> |  |  |  |  |  |  |
| N (%) | 57 (35) | 48 (30) | 33 (32) | 24 (26) | 23 (40) | 24 (35) |
| Uses pacifier |  |  |  |  |  |  |
| N (%) | 80 (49) | 86 (54) | 50 (49) | 42 (46) | 30 (53) | 43 (63) |
| Duration of breastfeeding (mo.)<sup>2</sup> |  |  |  |  |  |  |
| Mean (±SD) | 7.5 (±4.7) | 7.1 (±4.5) | 7.4 (±4.4) | 6.8 (±4.1) | 7.7 (±5.3) | 7.5 (±4.9) |
| Received ≥1 dose of pneumococcal conjugate vaccine |  |  |  |  |  |  |
| N (%) | 3 (2) | 4 (3) | 2 (2) | 4 (4) | 1 (2) | 0 (0) |
| Severe bulging |  |  |  |  |  |  |
| N (%) | 45 (28) | 41 (26) | 33 (32) | 30 (33) | 12 (21) | 10 (15) |
| Bilateral AOM |  |  |  |  |  |  |
| N (%) | 61/160 (38) | 68/158 (43) | 41/100 (41) | 34/90 (38) | 19/57 (33) | 34/67 (51) |
| Fever ≥39°C in previous 24 hours |  |  |  |  |  |  |
| N (%) | 24 (15) | 18 (11) | 16 (16) | 13 (14) | 8 (14) | 5 (7) |
| Pneumococcal carriage |  |  |  |  |  |  |
| Any, N (%) | 102/159 (64) | 91/159 (57) |  |  |  |  |
| PSSP, N (%) | 78/154 (52)<sup>3</sup> | 63/149 (45)<sup>3</sup> | 78/97 (81)<sup>3</sup> | 63/81 (78)<sup>3</sup> |  |  |
| PNSP, N (%) | 19/154 (13)<sup>3</sup> | 18/149 (13)<sup>3</sup> | 19/97 (19)<sup>3</sup> | 18/81 (22)<sup>3</sup> |  |  |

<sup>1</sup>No parents reported smoking inside the household.
<sup>2</sup>Duration of any breastfeeding at the time of the enrollment visit.
<sup>3</sup>Prevalence estimates are presented as median (95% confidence interval) based on the bootstrap approach (see **Methods** and **Text S1**) taken to correct for bias from incomplete testing of specimens.

### Impact of amoxicillin-clavulanate therapy for AOM against pneumococcal colonization

Among children from whom swabs were obtained as scheduled for the end-of-treatment visit, we detected pneumococcal carriage in 18 of 146 (12%) randomized to amoxicillin-clavulanate and 63 of 141 (45%) randomized to placebo (**Figure 2**), indicating 73% (95%CI: 58% to 84%) lower prevalence of pneumococcal colonization at end-of-treatment attributable to immediate antimicrobial therapy. Only 4% of children in the amoxicillin-clavulanate arm, compared to 34% in the placebo arm, carried PSSP at end-of-treatment. In contrast, 8% and 11% of children in the treatment and placebo arms, respectively, carried PNSP at end-of-treatment. These findings signified 88% (76% to 96%) and 25% (–54% to 65%) reductions in prevalence of PSSP and PNSP carriage at end-of-treatment due to immediate amoxicillin-clavulanate. In total, 32% (23% to 42%) prevalence of pneumococcal carriage among children treated for AOM could be averted by immediate amoxicillin-clavulanate therapy, almost entirely due to the prevention of PSSP carriage in 30% (21% to 38%) of children who would otherwise be expected to carry PSSP.

**Figure 2:**
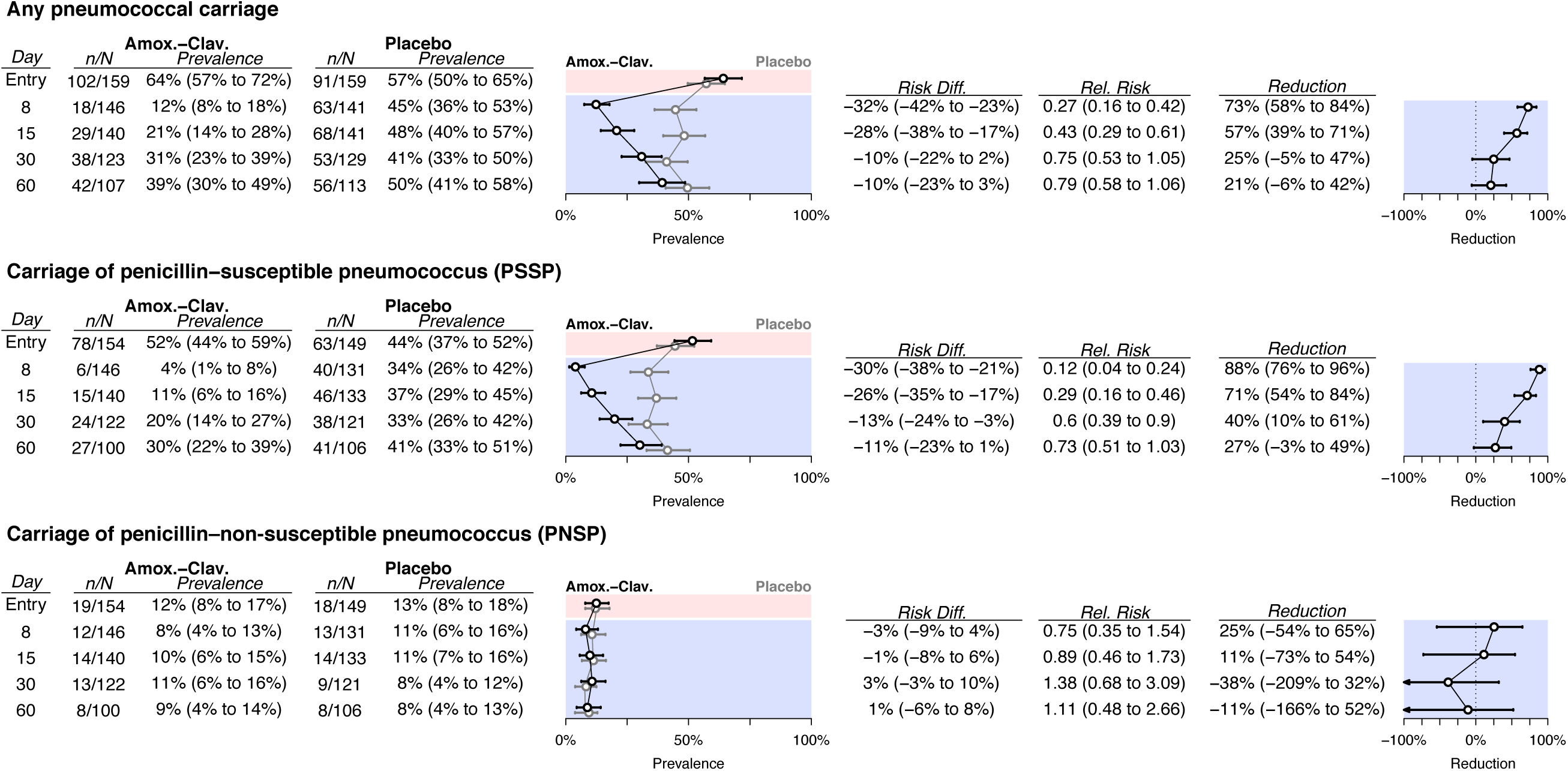
Impact of immediate amoxicillin-clavulanate on carriage of *S. pneumoniae* with varying susceptibility to penicillin. Tables and plots indicate the number carrying *S. pneumoniae*, PSSP, and PNSP, measured out of all swabs with microbiological determinations available (carriage presence/absence, or carriage presence/absence and penicillin susceptibility, given carriage). Prevalence estimates are presented as median (95% confidence interval) based on the bootstrap approach (see **Methods** and **Text S1**) taken to correct for bias from incomplete testing of specimens; consequently, median reported estimates may not equal *n*/*N*.

The magnitude of the impacts of immediate antimicrobial treatment on prevalence of any pneumococcal carriage and PSSP carriage declined over successive visits to 21% (–6% to 42%) and 27% (–3% to 49%), respectively, by 60d after treatment (**Figure 2**). We did not detect strong statistical evidence (defined by a 95%CI entirely above zero) that immediate amoxicillin-clavulanate impacted prevalence of PNSP carriage at any visit. Only 8% to 11% of children in either arm carried PNSP at any follow-up visit.

### Variation in outcomes according to carriage at enrollment

Stratifying analyses by carriage at the enrollment visit afforded a view into how antimicrobial therapy for AOM differentially impacted the clearance and acquisition of PSSP and PNSP colonization (**Figure 3** and **Table 2**). Among children carrying PSSP at enrollment, between 61% and 67% of those assigned placebo carried pneumococci at follow-up visits, whereas prevalence among children assigned antimicrobial therapy dropped to 7% by end-of-treatment and reached only 35% after 60d. This signified 89% (77% to 98%) lower prevalence of pneumococcal carriage at end-of-treatment due to immediate amoxicillin-clavulanate, with 45% (17% to 66%) lower prevalence in the treatment arm persisting up to 60d after treatment. No children who carried PSSP at enrollment were observed to carry PNSP at any follow-up visit (**Table 2**). In absolute terms, immediate amoxicillin-clavulanate prevented 54% (40% to 68%) of baseline PSSP carriers from carrying PSSP at end-of-treatment, relative to PSSP carriers assigned placebo; the averted prevalence of PSSP carriage as of 60d after treatment amounted to 28% (9% to 48%).

**Table 2:**
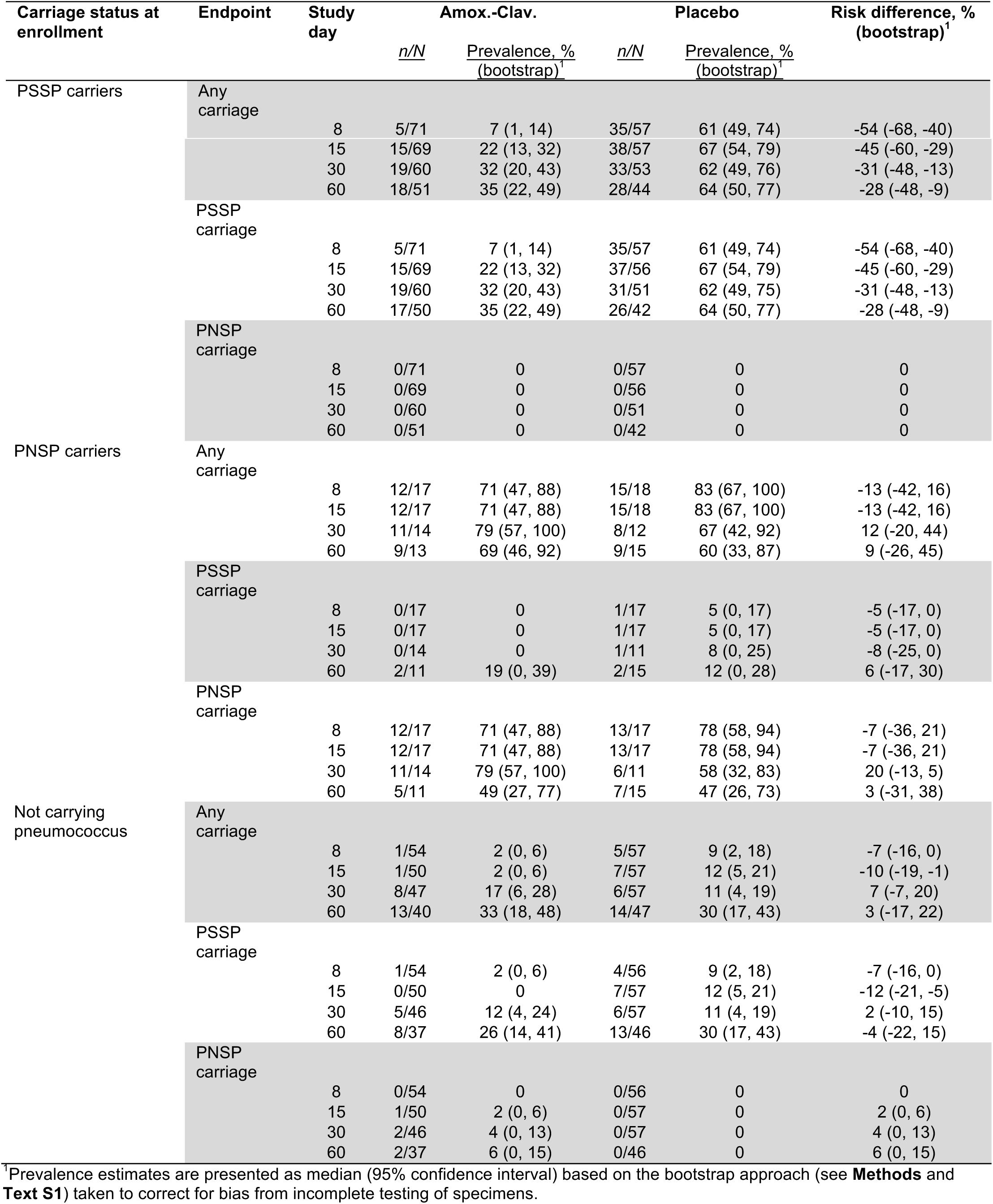
Prevalence of pneumococcal carriage averted by immediate antimicrobial therapy, according to carriage status at enrollment.

**Figure 3:**
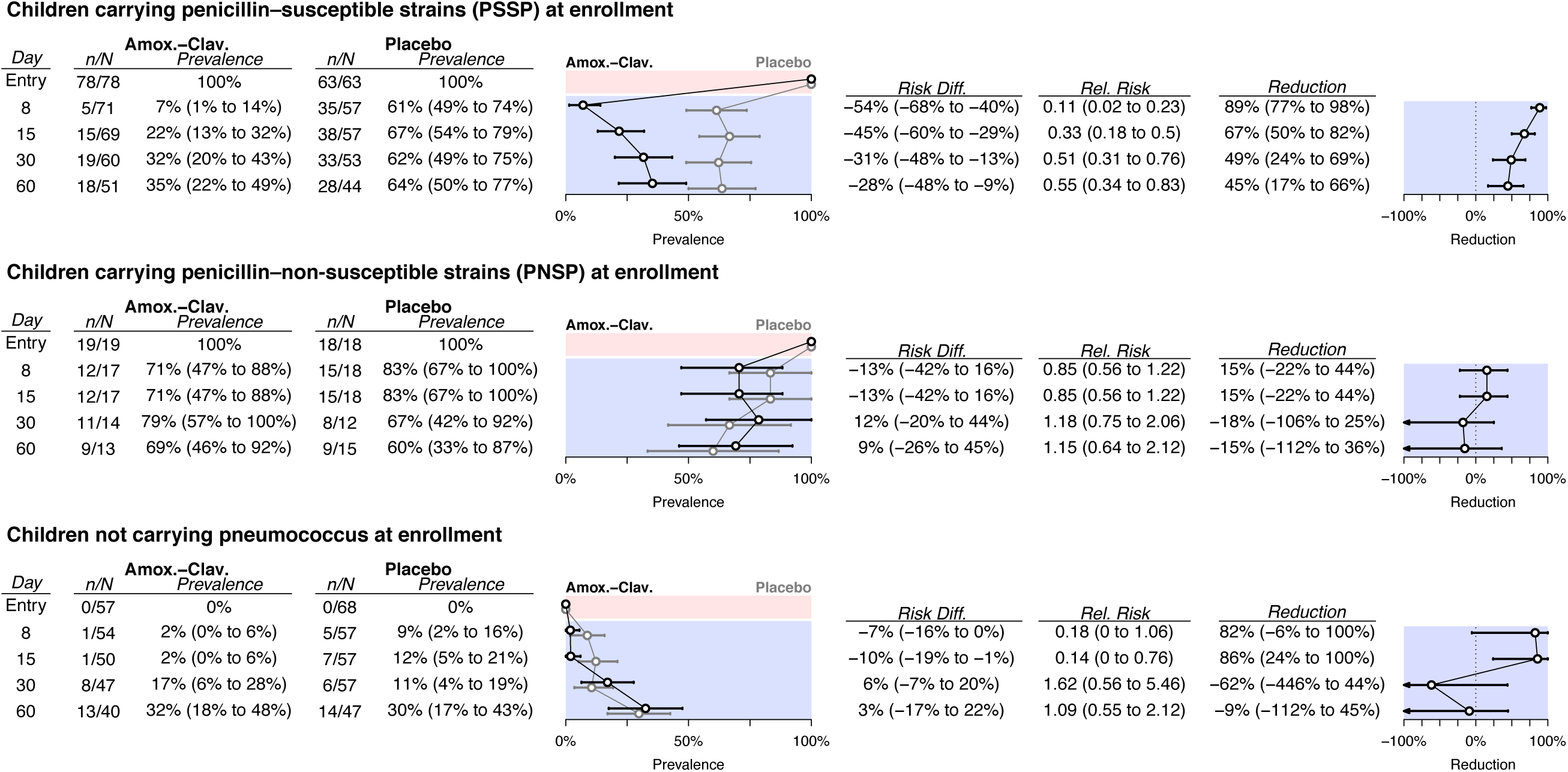
Impact of immediate amoxicillin-clavulanate on carriage of *S. pneumoniae*, according to carriage at enrollment. Tables and plots indicate the number carrying *S. pneumoniae* at follow-up visits, according to carriage at enrollment, measured out of all swabs with microbiological determinations available (carriage presence/absence, or carriage presence/absence and penicillin susceptibility, given carriage); prevalence estimates are presented as median (95% confidence interval) based on the bootstrap approach (see **Methods** and **Text S1**) taken to correct for bias from incomplete testing of specimens. We expand on results presented in this Figure—detailing susceptibility of isolates carried at follow-up—in **Table 2**.

Among children carrying PNSP at enrollment, 71% and 83% carried pneumococcus at end-of-treatment after randomization to amoxicillin-clavulanate and placebo, respectively, indicating 15% (–22% to 44%) lower colonization prevalence due to treatment (**Figure 3**). Impact estimates at subsequent follow-up visits remained statistically indistinguishable from zero. By 60d after treatment, 69% and 60% of children assigned immediate amoxicillin-clavulanate and placebo, respectively, carried *S. pneumoniae*. Of the children carrying PNSP at enrollment and assigned amoxicillin-clavulanate, none carried PSSP until 60d after treatment (**Table 2**), whereas 5% to 8% prevalence of PSSP carriage was detected at follow-up visits by children assigned placebo.

Among children who were not colonized at enrollment, we estimated that immediate antimicrobial therapy resulted in 82% (–6% to 100%) and 86% (24% to 100%) lower prevalence of pneumococcal colonization at end-of-treatment and 15d after treatment, respectively (**Figure 3**). However, prevalence of colonization among such children was low; the averted prevalence 15d after treatment amounted to only 10% (1% to 19%). We did not identify statistical evidence that treatment impacted carriage prevalence at subsequent follow-up visits. The majority of pneumococcal isolates collected after treatment from children who were not colonized at enrollment were penicillin-susceptible (**Table 2**). Among children who were not colonized at enrollment, no PNSP carriage was detected among those assigned placebo, whereas PNSP prevalence reached 6% (0% to 15%) by 60d after treatment among those assigned antimicrobial therapy. However, this reflected acquisition of PNSP by only three children, with zero to two children simultaneously carrying PNSP at any scheduled visit.

### Selective impact of amoxicillin-clavulanate therapy for AOM

Owing to the differential impacts of antimicrobial therapy on clearance and acquisition of PSSP and PNSP, the proportion of carried strains showing reduced susceptibility to penicillin at end-of-treatment was 2.74 (95%CI: 1.62 to 4.88) times higher among children assigned immediate amoxicillin-clavulanate (12 of 18, 67%) than among children assigned placebo (13 of 53, 25%; **Table 3**). This effect persisted at least 30d after treatment, at which point the proportion of carried strains showing reduced susceptibility to penicillin was 2.28 (1.18 to 5.53) times higher among children assigned immediate antimicrobial therapy (35%) than among those assigned placebo (15%). By 60d after treatment, the proportion of pneumococcal isolates with reduced susceptibility to penicillin remained higher among children assigned antimicrobial therapy (cRR 1.40 (0.64 to 3.20)), although this difference no longer reached conventional thresholds for statistical significance.

**Table 3:** Conditional risk ratios for carried *S. pneumoniae* to show reduced susceptibility to penicillin, given immediate antimicrobial therapy.

| Study day | Amox.-Clav. |  | Placebo |  | Conditional risk ratio |
| --- | --- | --- | --- | --- | --- |
|  | n/N | Proportion, % | n/N | Proportion, % |  |
| 8 | 12/18 | 67 (45, 87) | 13/53 | 25 (15, 35) | **2.74 (1.62, 4.88)** |
| 15 | 14/29 | 48 (32, 65) | 14/60 | 23 (14, 33) | **2.08 (1.22, 3.69)** |
| 30 | 13/37 | 35 (22, 48) | 7/45 | 15 (7, 25) | **2.28 (1.18, 5.53)** |
| 60 | 8/35 | 23 (12, 34) | 8/49 | 16 (8, 25) | 1.40 (0.64, 3.20) |

### Penicillin-resistant pneumococcal carriage

Three children carried *S. pneumoniae* strains that were classed as fully resistant to penicillin (defined by an oxacillin disk reading <20mm and MIC>2µg/mL) (**Figure 4**). Two of these children carried strains registering an intermediate MIC at enrollment, and were found to carry penicillin-resistant isolates of the same serotype at least once after treatment with amoxicillin-clavulanate as the study drug or rescue therapy. A third child who received amoxicillin-clavulanate carried a strain registering a resistant MIC at all study visits, with the exception of the end-of-treatment visit, at which an intermediate MIC was detected. We detail the clinical course of these children in the supplemental text (**Text S2**).

**Figure 4:**
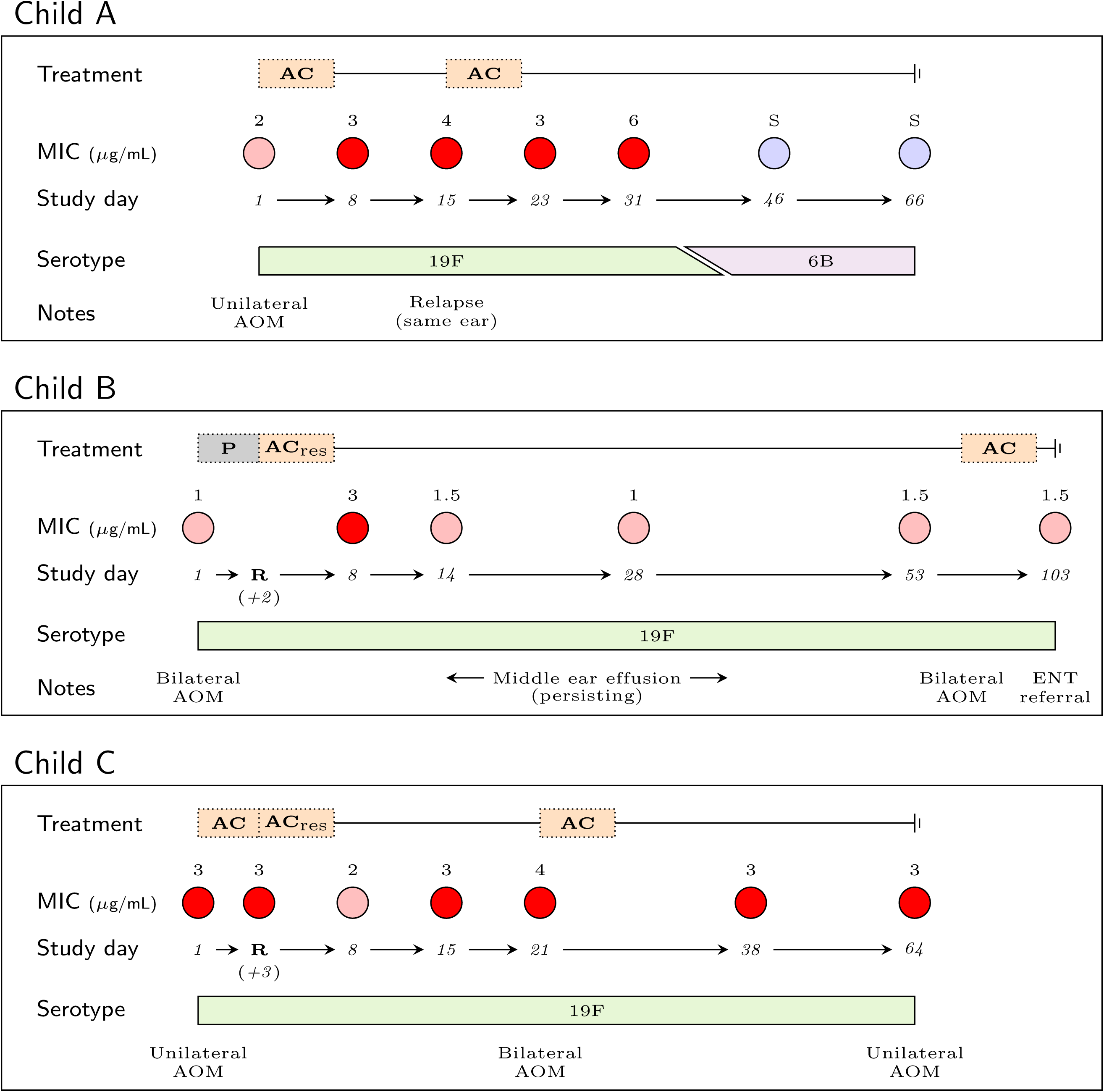
Observations among children who carried resistant isolates during the study. Study drug and rescue therapy assignments (P: placebo; AC: amoxicillin-clavulanate) are illustrated together with isolates (with MIC values) taken from children from the enrollment visit to the last visit performed. Study days are numbered from the first day of treatment with study drug or rescue therapy. Colors indicate penicillin sensitivity of the carried strain: blue, penicillin sensitive; pink, intermediate penicillin resistance; red, penicillin resistance.

## DISCUSSION

We report a sustained, selective impact of antimicrobial treatment for AOM on pneumococcal carriage among young children in a high-income country. In this trial, children randomized to immediate amoxicillin-clavulanate therapy were less likely than those randomized to placebo to carry PSSP up to 60d after treatment. By end-of-treatment, PNSP constituted 67% of carried strains among children assigned antimicrobial therapy, compared to 25% of carried strains among children assigned placebo; differences between the treatment and placebo arms in the susceptibility of carried strains persisted at least 30d after treatment. Despite this, antimicrobial treatment did not impact absolute prevalence of PNSP carriage. These findings clarify the magnitude and duration of the impact of immediate antimicrobial therapy, as compared to watchful waiting, on carriage of *S. pneumoniae* with varying susceptibility to penicillin.

In this trial, repeated sampling of nasopharyngeal carriage at enrollment and follow-up visits illustrated the epidemiological basis of selection under antimicrobial treatment through the differential clearance and acquisition of PSSP and PNSP. Under circumstances resembling watchful waiting, 39% of PSSP carriers initially assigned placebo cleared colonization by end-of-treatment, either naturally or due to receipt of antimicrobial drugs as rescue therapy, compared to 18% of PNSP carriers. Immediate antimicrobial therapy cleared PSSP colonization by end-of-treatment in an additional 54% of children, but had no measurable impact on colonization among PNSP carriers. Among children who were not colonized at enrollment, immediate antimicrobial therapy lowered the risk of acquiring PSSP by >80% over the first two weeks after treatment. These measurements of the acquisition and clearance of susceptible and non-susceptible pneumococcal lineages under differing treatment protocols can inform transmission models, guiding the assessment of population-level consequences of differing antimicrobial prescribing strategies.

Among placebo-controlled trials of antimicrobial treatment for AOM, this study enrolled the largest sample of children and included the longest continuous monitoring of carriage [16]. By stratifying according to carriage status at enrollment, our analysis is the first to assess effects of antimicrobial treatment for AOM on clearance and acquisition of PSSP and PNSP. Several other trials of antimicrobial therapy for AOM in high-income countries provide complementary insights. Consistent with our findings, no difference in absolute prevalence of PNSP was reported at end-of-treatment or 21-25d after treatment among children assigned 10d of high-dose amoxicillin-clavulanate compared to those assigned placebo [3], or at the end of a 10d versus 5d course of high-dose amoxicillin-clavulanate [9]. In both trials, the proportions of pneumococcal isolates showing reduced susceptibility to penicillin at end-of-treatment were higher among children assigned 10d amoxicillin-clavulanate than among children assigned either placebo or 5d amoxicillin-clavulanate. This pattern suggests selection was mediated by reductions in PSSP persistence or acquisition, consistent with our observations [3,9]. Similar changes in the proportion of isolates showing reduced susceptibility were evident after treatment in a trial comparing immediate antimicrobial prescribing and watchful waiting [10], although effects on absolute prevalence of PSSP and PNSP were not reported, and carriage was assessed only 12d after treatment. Unified reporting of absolute and conditional effects on bacterial carriage and susceptibility will facilitate direct comparison of results from future trials [11]. Our finding that reductions in PSSP carriage persist up to 60d after treatment also suggests that future studies should monitor carriage over longer follow-up windows than previous trials have included.

Whereas we did not detect increases in PNSP carriage prevalence due to antimicrobial treatment, this observation has been reported in several studies undertaken in lower-income countries with higher PNSP prevalence. In the Dominican Republic, children with respiratory tract infections randomized to a low-dose, 10d amoxicillin regimen had higher prevalence of PNSP carriage after treatment than children randomized to a high-dose, 5d regimen [17]. In Malawi, increases in PNSP carriage prevalence were reported after treatment of malaria with sulfadoxine-pyrimethamine and treatment of suspected bacterial infections with cotrimoxazole [18]. Whereas most studies of antimicrobial therapy for AOM have been conducted in the United States and Europe [16], these findings suggest trials or observational studies are warranted in low-and middle-income countries to guide appropriate treatment practices for such settings, as higher PNSP prevalence may impact the relationship between antimicrobial treatment and carriage dynamics.

It is important to note that our analysis measured only the direct effects of antimicrobial therapy on carriage dynamics [19]. Large-scale changes in treatment practices may spur indirect effects on carriage of PSSP and PNSP among children not treated for AOM. Several lines of evidence demonstrate indirect effects of antimicrobial treatment interventions on pneumococcal carriage and susceptibility. Mass azithromycin administration for the prevention of trachoma caused community-wide increases in prevalence of macrolide-resistant *S. pneumoniae* carriage in a cluster-randomized trial in Ethiopia [20]. In ecological studies in higher-income settings, outpatient antibiotic prescribing predicts the proportion of pneumococcal isolates showing reduced penicillin susceptibility at geographic scales ranging from municipalities to countries [21,22].

This study has several limitations. It is important to note that the study treatment (40mg of amoxicillin plus 5.7mg of clavulanate per kg of body weight daily) differs from the high-dose amoxicillin-clavulanate regimen recommended in the US (90mg of amoxicillin plus 6.4mg clavulanate per kg of body weight daily). Notably, high-dose amoxicillin-clavulate did not deliver larger reductions in carriage or lower rates of clinical failure in the US than 40mg/5.7mg/kg amoxicillin-clavulanate delivered in this trial in Finland [3]. Because our trial was designed to assess clinical failure as a primary endpoint, stratified analyses addressing carriage of PNSP, and outcomes among children who carried PNSP at enrollment, had limited statistical power. Although data from four time points up to 60d after treatment provided a more detailed view of pneumococcal carriage dynamics than other trials have afforded [3,9,10], swabs were not obtained from 22% and 32% of children from follow-up visits 30d and 60d after treatment, respectively. While our analysis addresses treatment effects on *S. pneumoniae*, effects on other pathogens and commensal flora within the microbiome are also of importance [23]. Last, our analysis does not address the association of carriage endpoints with antimicrobial prescribing for secondary AOM episodes or other causes during follow-up. An advantage of the modified intention-to-treat framework we employ is that this circumstance does not hinder our interpretation of carriage outcomes as causal effects of the initial randomized assignment.

This trial was undertaken before the introduction of pneumococcal conjugate vaccination in the Finnish national immunization program. Importantly, our stratification of analyses by the penicillin susceptibility of colonizing lineages provides a basis for transporting effect size estimates, as treatment effects in other populations may be predicted by the distribution of penicillin susceptibility among colonizing serotypes. Whereas a higher proportion of vaccine-targeted serotypes than non-vaccine serotypes tended to be resistant to antimicrobial drugs in the pre-vaccine era, increases in the proportion of resistant non-vaccine serotypes have been reported amid serotype replacement, generally mitigating any net change in the prevalence of resistance [24–26].

In conclusion, we find that immediate amoxicillin-clavulanate therapy for AOM facilitates clearance and delays acquisition of PSSP in the nasopharynx, indirectly increasing the proportion of carried pneumococcal strains showing reduced susceptibility to penicillin. We detected no increase in absolute prevalence of PNSP among children assigned immediate treatment as compared to children assigned placebo. These findings suggest antimicrobial treatment for AOM may not increase risk for individual children to carry penicillin–non-susceptible strains of *S. pneumoniae*, although selective impacts of widespread treatment of AOM remain of concern at the population level. Quantifying the effects of different AOM treatment strategies on carriage of PSSP and PNSP provides a basis for evaluating antimicrobial resistance risks against the clinical benefits of antimicrobial therapy for AOM.

## SUPPORTING INFORMATION CAPTIONS

**S1 Text. Bootstrap approach to statistical inference and missing-data bias correction.** We detail the approach taken to statistical inference and prevalence estimation in the presence of missing data from laboratory results.

**S2 Text. Clinical course of children carrying fully penicillin-resistant *S. pneumoniae*.** We describe the clinical course of carriage and disease in children who carried fully-resistant strains.

**S1 Table. Baseline attributes of children with missing and complete carriage data.** We show that children with complete or missing data from scheduled follow-up visits did not differ in baseline characteristics.

## Text S1 Bootstrap approach to statistical inference and missing-data bias correction

We used the bootstrap to correct for missing-data biases under incomplete testing in prevalence estimates of PSSP and PNSP. Pneumococcal carriage prevalence and prevalence of PSSP and PNSP at each follow-up visit were subject to differential missingness under incomplete testing of specimens for pneumococcus, and under incomplete testing for resistance in specimens that tested positive for carriage. This could lead to bias under the conditions laid out below:

For arm *A*, define *N*_*A*_ as the total number of children, *C*_*A*_=*p*_*C|A*_*N*_*A*_ as the total number of carriers, and *S*_*A*_=*p*_*S|A*_*N*_*A*_=*p*_*S|CA*_*C*_*A*_ as the total number of carriers of pneumococcus with susceptibility *s*. Define *t*_*C|A*_ and *t*_*S|CA*_, respectively, as the proportions of children tested for pneumococcal carriage and the proportion of carriage-positive specimens for which resistance was characterized. Under an assumption that missingness is unrelated to carriage status within any study arm, excluding untested specimens does not lead to bias in naive measurements of pneumococcal carriage prevalence

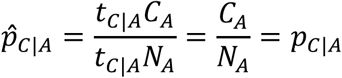

or in measurements of the proportion of carriage-positive specimens with susceptibility *s*:

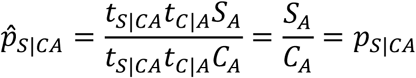

However, measures of the prevalence of pneumococcus with susceptibility *s* were subject to bias when resistance data were missing:

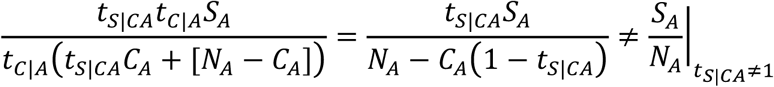

We therefore measured the prevalence of pneumococci with susceptibility *s* from the unbiased measures 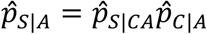 using the bootstrap-resampled estimates of 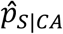 and 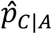.

## Text S2 Clinical course of children carrying fully penicillin-resistant *S. pneumoniae*

*“Child A”* carried an intermediately-resistant (MIC=2µg/mL) strain (serotype 19F) at enrollment and was assigned amoxicillin-clavulanate. The child had unilateral AOM at the enrollment visit, and at the end-of-treatment visit (day 8) his symptoms and otoscopic signs were improving. The child had a relapse of AOM in the same ear as the initial episode on day 15, and was prescribed amoxicillin-clavulanate for 7 days. Both ears were diagnosed healthy at the end of the second course of antimicrobial treatment on day 23. No other episodes of AOM were diagnosed during the follow-up. The strain of *S. pneumoniae* was found to be resistant from end-of-treatment to day 31, when the highest level of resistance was detected (MIC=6µg/mL). The child carried a penicillin-susceptible strain (serotype 6B) at subsequent visits (days 46 and 66).

*“Child B”* carried an intermediately-resistant (MIC=1µg/mL) strain (serotype 19F) at enrollment and was randomized to placebo. At enrollment, she had bilateral AOM. During follow-up the symtoms and otoscopic signs deteriorated, and rescue treatment was initiated on day 3. After that, the child’s overall condition and otoscopic signs improved, but she still had middle ear effusion in both ears until day 97, when she was diagnosed with bilateral AOM and received amoxicillin-clavulanate for 7 days. The carried 19F strain registered a resistant MIC (3µg/mL) at end-of-treatment, but intermediate resistance (MIC 1 to 1.5µg/mL) at subsequent visits to day 103. Eventually, the child was sent for referral to an ear, nose and throat specialist due to persistent middle ear effusion.

*“Child C”* presented with unilateral AOM and carried a resistant (MIC=3µg/mL) strain (serotype 19F) at enrollment, and received amoxicillin-clavulanate. On day 4 her overall condition had worsened with bulging of the tympanic membrane in both ears. As the study personnel remained blinded to the study group assignment until the completion of the whole trial, she received rescue treatment (amoxicillin-clavulanate) for additional 7 days. On day 15 both ears were healthy. On day 21 she had a bilateral episode of AOM and was prescribed amoxicillin-clavulanate for 7 days. On day 38 both ears were healthy again. On day 64 she had unilateral AOM that was treated with watchful waiting. The carried strain showed resistant MIC readings at all visits except the end-of-treatment visit, when borderline intermediate resistance was detected (MIC=2µg/mL).

**Table S1.** Baseline attributes of children with missing and complete carriage data.

| Characteristic | Amox.-Clav. |  | Placebo |  |
| --- | --- | --- | --- | --- |
|  | Children missing carriage or resistance data at any visit (N=80) | Children with no missing data at any visit (N=82) | Children missing carriage or resistance data at any visit (N=79) | Children with no missing data at any visit (N=81) |
| Age (mo.) |  |  |  |  |
| Mean (±SD) | 16.5 (±7.5) | 14.6 (±6.9) | 16.7 (±8.0) | 15.4 (±7.9) |
| Male child |  |  |  |  |
| N (%) | 43 (54) | 49 (60) | 41 (52) | 50 (62) |
| Received antimicrobial drugs within preceding month |  |  |  |  |
| N (%) | 20 (25) | 17 (21) | 13 (16) | 21 (26) |
| Number of siblings |  |  |  |  |
| 0, N (%) | 41 (51) | 32 (39) | 35 (44) | 31 (38) |
| 1–2, N (%) | 36 (45) | 43 (52) | 43 (54) | 46 (57) |
| 3 or more, N (%) | 3 (4) | 7 (9) | 1 (1) | 4 (5) |
| Daycare attendance |  |  |  |  |
| N (%) | 47 (59) | 40 (49) | 49 (62) | 38 (47) |
| Parental smoking<sup>1</sup> |  |  |  |  |
| N (%) | 29 (36) | 28 (35) | 22 (28) | 26 (32) |
| Uses pacifier |  |  |  |  |
| N (%) | 39 (49) | 41 (50) | 44 (56) | 42 (52) |
| Duration of breastfeeding (mo.)<sup>2</sup> |  |  |  |  |
| Mean (±SD) | 7.8 (±4.2) | 7.3 (±5.2) | 6.5 (±4.2) | 7.7 (±4.7) |
| Received ≥1 dose of pneumococcal conjugate vaccine | | | | |
| N (%) | 1 (1) | 2 (2) | 1 (1) | 3 (4) |
| Severe bulging |  |  |  |  |
| N (%) | 25 (31) | 20 (24) | 25 (32) | 16 (20) |
| Bilateral AOM |  |  |  |  |
| N (%) | 32 (41) | 29 (36) | 34 (44) | 34 (42) |
| Fever ≥39°C in previous 24 hours | | | | |
| N (%) | 13 (16) | 11 (13) | 11 (14) | 7 (9) |
| Pneumococcal carriage |  |  |  |  |
| Any, N (%) | 51 (66) | 51 (62) | 48 (62) | 43 (53) |
| PSSP, N (%) | 36 (50) | 42 (51) | 29 (43) | 34 (42) |
| PNSP, N (%) | 10 (14) | 9 (11) | 9 (13) | 9 (11) |

<sup>1</sup>No parents reported smoking inside the household.
<sup>2</sup>Duration of any breastfeeding at the time of the enrollment visit.

